# The Interaction Between The Noradrenergic Neurons In The Locus Coeruleus And Beta-1 Adrenergic Receptor (β1-AR) In The Cardiomyocytes Was Implicated In Modulating SUDEP

**DOI:** 10.1101/2022.01.10.475630

**Authors:** XiTing Lian, Qian Yu, HaiXiang Ma, LeYuan Gu, Qing Xu, YuLing Wang, Yue Shen, HongHai Zhang

## Abstract

Sudden unexpected death of epilepsy (SUDEP) is the leading cause of of death in patients with epilepsy. Due to the complicated pathogenesis of SUDEP, however, the exact mechanism of SUDEP remains elusive. Currently, although it is recognized that the seizure-induced respiratory arrest (S-IRA) may be a main cause for SUDEP, other factors resulting in SUDEP can not be excluded e.g arrhythmias. Our previous findings indicated that the incidence of S-IRA and SUDEP evoked by acoustic stimulation or pentetrazol (PTZ) injection was significantly reduced by atomoxetine, a norepinephrine reuptake inhibitor (NRI), suggesting that noradrenergic neurotransmission modulates S-IRA and SUDEP. Given that norepinephrine acts on the target to modulate respiratory and circulation function by targeting adrenergic receptor α and beta (a-AR and β-AR) and the arrhythmias can be contributed to SUDEP. Meanwhile, to further test whether cardiac factors are implicated in S-IRA and SUDEP, we choose esmolol hydrochloride, a selective antagonist of β1-AR to test it in our models. Our findings demonstrated that the lower incidence of S-IRA and SUDEP evoked by acoustic stimulation or PTZ injection in DBA/1 mice by administration with atomoxetine was significantly reversed by intraperitoneal (IP) of esmolol hydrochloride. Importantly, the data of electrocardiogram (ECG) showed that the cardiac arrhythmia including the ventricular tachycardia, ventricular premature beat and atrioventricular block can be evoked by acoustic stimulation or PTZ injection in our model. Administration of atomoxetine significantly reduced these arrhythmias and the incidence of S-IRA and SUDEP in our models. However, administration of esmolol hydrochloride with the dose without affecting ECG and mortality changing of DBA/1 significantly blocking the protective effects of atomoxetine on S-IRA and SUDEP in our models. Thus, the dysfunction of respiratory and circulation may be implicated in the pathogenesis of S-IRA and SUDEP. Enhancing the central norepinephrinergic neurotransmission in the brain contributes to inhibition of seizure-induced respiratory arrest by targeting β1-AR locating in the cardiomyocytes. Furthermore, the suppression effects of S-IRA by atomoxetine was significantly reversed by the norepinephrine neuronal degradation in the LC in our models. Furthermore, PTZ-induced Tyrosine hydroxylase (TH), the rate-limiting enzyme in the synthesis of norepinephrine, activity but not TH content from the serum of left ventricle and the whole heart tissue was reduced following the S-IRA. Our findings will show a new light on decoding the pathogenesis of SUDEP concerning the pathway between the LC and heart.

## 1. INTRODUCTION

Sudden unexpected death in epilepsy (SUDEP) is the most severe complication leading to the mortality of epilepsy patients, especially those with anti-epileptic drug resistance^1–4^. Although some important advancements concerning the risk factors, biomarkers and pathogenesis of SUDEP had been obtained, to our certain knowledge, decoding SUDEP seems to be a huge challenge in views of the complicated mechanism of SUDEP. Indeed, seizure-induced respiratory arrest (S-IRA) is an important cause resulting in the death of epilepsy including our previous findings^5, 6^. However, the underlying mechanism of SUDEP is complex, which is not caused by a single factor, but the result of the comprehensive action of multiple factors^7, 8^.

Previous findings showed that the cardiac and respiratory function were simultaneously suppressed to induce the S-IRA and SUDEP evoked by acoustic stimulation in DBA/1 mice, suggesting that the autonomic nerve including the sympathetic and parasympathetic nerve were involved in the course of S-IRA and SUDEP^9^. Our previous research indicates that atomoxetine, a norepinephrine reuptake inhibitor, can significantly reduce the incidence of SUDEP in DBA/1 mice by increasing the content of NE in the brain^5, 10^. It was well known that norepinephrine transmission is closely related to the activity of sympathetic neurons and modulates the respiratory and circulation function by targeting the adrenergic α1, α2, β1, and β2 receptors. The findings from our and other laboratories showed α1 and α2 receptors were implicated in the S-IRA and SUDEP medicated by atomoxetine in the DBA/1 mice models^11, 12^. As for β1 and β2 receptors, especially β1 receptor, the postganglionic nerve endings of cardiac sympathetic neurons mainly release NE and act on β1receptors which locate in the myocardial cell and play a key role in modulating the cardiac function^13, 14^. Therefore, we choose esmolol hydrochloride, a selective antagonist of β1 adrenergic receptor (β1-AR) to test whether the β1-AR can reverse the lower incidence of S-IRA and SUDEP mediated by atomoxetine to test the interaction between the brain and heart to modulate SUDEP. Based on our previous research, enhancing the synaptic transmission efficiency of NE in the brain can effectively reduce the incidence of SUDEP^5^. Given this, we can’t exclude the occurrence of SUDEP may be the result of the regulation of the central and peripheral sympathetic nerve. Studies have shown that the locus coeruleus (LC) is the main nucleus for the synthesis and release of NE in the brain^15^. In order to better explore the relationship between the content of NE in the brain, especially in the LC and SUDEP, we choose the N-(2-chloroethyl)-N-ethyl-2-bromoben-zylamine hydrochloride (DSP-4), a selective neurotoxin for the locus coeruleus noradrenergic system in the rodent brain, to destroy the nerve terminal^16^, which results in a rapid, transient reduction in NE and NE transporter in many brain regions receiving variable innervation from the LC in our models^17^. Meanwhile, the previous study showed that the noradrenergic nerve fibers of the LC projected into the nucleus tractus solitarius (NTS), a circulatory regulation center, the latter projected the sympathetic neurons fibers into the stellate ganglion (SG) by the long distance, which composed the cardiac sympathetic preganglionic fibers to release the NE to combine the β1-AR in the cardiomyocyte membrane. Therefore, it may be the possible interaction between the LC and heart to modulate SUDEP in our models.

Based on this, we observed the effects of administration of DSP-4 to reduce the concentration of NE in LC and esmolol to block the β1-AR to test whether the lower incidence of S-IRA and SUDEP by atomoxetine can be reversed, respectively. It turns out that the suppression of S-IRA and SUDEP by atomoxetine can be significantly reversed by administration of DSP-4 and esmolol, separately, which highly suggests the sympathetic excitability of brain and heart have similar effect on preventing SUDEP. Although we did not verify the anatomical construction and function of each specific nucleus relating to sympathetic axis between the brain and heart, the results of our experiment shed an important light on the central peripheral sympathetic axis in mediating the occurrence of SUDEP and this reminds us when treating a patient with epilepsy, adrenergic receptor blocker may be a risk factor to cause the death of patient. Furthermore, PTZ-induced TH, the rate-limiting enzyme in the synthesis of norepinephrine, activity but not TH content from the serum of left ventricle and the whole heart tissue was reduced following the S-IRA. What’s more, our preliminary findings demonstrated that the interaction between the noradrenergic neurons in the LC and β1-AR in the cardiomyocytes was implicated in modulating SUDEP and modulating the interaction will be a potential target to intervene for preventing SUDEP.

## 2. MATERIALS AND METHODS

### 2.1 Animals

All experimental procedures were the agreement with the National Institutes of Health Guidelines for the Care and Use of Laboratory Animals and approved by the Animal Advisory Committee of Zhejiang University. DBA/1 mice were housed and bred in the Animal Center of Zhejiang University School of Medicine and provided with rodent food and water ad libitum. DBA/1 mice of either sex were used in this experiment, as previous studies have shown, gender is not a variable that affects the S-IRA of DBA/1 mice^18^. Aiming at acoustic stimulation murine model, DBA/1 mice were “primed” starting from postnatal day 26-28 to establish the consistent susceptibility to audiogenic seizures and S-IRA as previously described^5, 10, 19, 20^. For the PTZ-evoked seizure model, PTZ was administered to non-primed DBA/1 mice at approximately 8 weeks of age.

### 2.2 Seizure induction and resuscitation

S-IRA was established by acoustic stimulation as we previously described^5, 10, 19, 20^. For the acoustic stimulation model, each mouse was placed in a cylindrical plexiglass chamber in a sound-isolated room, and audiogenic seizures (AGSZs) were evoked by using an electric bell (96 dB SPL, Zhejiang People’s Electronics, China). Acoustic stimulation was given for a maximum duration of 60s or until the onset of tonic seizures and S-IRA in most mice in each group. S-IRA was evoked in all non-primed DBA/1 mice by a single dose IP administration of PTZ (Cat # P6500; Sigma-Aldrich,St. Louis, MO,) at 75mg/kg. Mice with S-IRA have resuscitated within 5s post the final respiratory gasp by a rodent respirator (YuYan Instruments, Shanghai, China).

### 2.3 Pharmacology experiment

#### 2.3.1 Effects of intraperitoneal administration of Esmolol on the atomoxetine-mediated suppression of S-IRA evoked by acoustic stimulation

The susceptibility of DBA/1 mice to S-IRA was confirmed 24 hours before atomoxetine or vehicle administration. To IP administration, atomoxetine and esmolol were dissolved in saline. Different dosages of esmolol were applied to vehicle control group and treatment groups, in which DBA/1 mice would be acoustically stimulated later. Atomoxetine (15 mg/kg, Ca # Y0001586; Sigma-Aldrich) or vehicle was intraperitoneal (IP) injection in DBA/1 mice 120min prior to acoustic stimulation,115min after infusion of atomoxetine (or vehicle), IP injection of esmolol (25mg/kg, 50mg/kg, H19991058; Qilu Pharmaceutical Co., Ltd.) or vehicle 5min before acoustic stimulation (n=5-8/per group). The incidence of S-IRA, latency to AGSZs, duration of wild running, clonic seizures, tonic-clonic seizures, and seizure scores were videotaped for offline analysis as described previously^21–23^. To exclude the effects of esmolol on the incidence of the mortality of DBA/1 mice, the group, subjected to confirmation of S-IRA before 24 hours to experiment, was pre-treated with the esmolol (50mg/kg, i.p) without administration of atomoxetine and acoustic stimulation in the same manner to observe the incidence of the mortality of DBA/1 mice and heart rate changes (n= 6). DBA/1 mice from the different pre-treatment groups, the monitoring of ECG was performed before acoustic stimulation and post-S-IRA.

#### 2.3.2 Effects of intraperitoneal administration of Esmolol on the atomoxetine-mediated suppression of S-IRA evoked by PTZ

A vehicle control group and treatment groups with atomoxetine and esmolol were included in testing of S-IRA in PTZ model. Atomoxetine (15 mg/kg) or vehicle was administered IP in DBA/1 mice 2h prior to IP injection of PTZ (75 mg/kg). Esmolol (50 mg/kg) or vehicle was given by IP infection 10min post administration of PTZ (n = 5-8/per group). The incidence of S-IRA, latency to GSZs, duration of wild running, clonic seizures, tonic-clonic seizures, and seizure scores were videotaped for offline analysis as described previously^21–23^. DBA/1 mice from the different pre-treatment groups, the monitoring of ECG was performed before and post IP administration of PTZ.

#### 2.3.3 Effects of intraperitoneal administration of DSP-4 on the atomoxetine-mediated suppression of S-IRA evoked by acoustic stimulation

Before a drug or vehicle treatment 24h, the susceptibility of primed DBA/1 mice to S-IRA was confirmed. To identify the effect of atomoxetine on S-IRA, N-(2-Chloroethyl)-N-ethyl-2-bromobenzylamin hydrochloride (DSP-4, C8417, Sigma-Aldrich) or vehicle for 50mg/kg intraperitoneally administered 1, 3 or 7 days prior to acoustic stimulation, and acoustic stimulation was given 2h after atomoxetine (15 mg/kg) IP injection (n=5-10/per group). The incidence of S-IRA, latency to AGSZs, duration of wild running, clonic seizures, tonic-clonic seizures, and seizure scores were videotaped for offline analysis as described previously^21–23^. Toexplore the effects of DSP-4 on noradrenergic neurons in the LC (locus coeruleus) of DBA/1 mice, the group was pre-treated with DSP-4 (50mg/kg, IP) or vehicle without administration of atomoxetine and acoustic stimulation. DBA/1 mice were sacrificed and perfused in 1, 3 or 7 days respectively to count the TH+ cells (n=3, n=5). TH+ cells were counted in 5 sections (n=5) for each animal, and compare DSP-4 treated with vehicle treated mice, n=3).

#### 2.3.4 Effects of microinjection of DSP-4 in the bilateral locus coeruleus (LC) on the atomoxetine-mediated suppression of S-IRA evoked by PTZ

DBA/1 mice with bilateral locus coeruleus guide cannula (62070, RWD Life Science Inc.,China) implantation (AP ± 0.9mm; ML - 5.45 mm; DV - 3.15 mm) for 1week will be used. 7 days before the induction of S-IRA, DSP-4 (200nl, 10µg/µL) or vehicle was separately administered by guide cannula to the every unilateral LC (n=8-9/per group). Atomoxetine or vehicle (15mg/kg,IP) was administered 120 mins prior to PTZ (75mg/kg, IP). The incidence of S-IRA, latency to GSZs, duration of wild running, clonic seizures, tonic-clonic seizures, and seizure scoreswere videotaped for offline analysis as described previously^21–23^. To explore the effects of microinjection of DSP-4 in the bilateral locus coeruleus on DBA/1mice noradrenergic neurons, these mice were sacrificed and perfused after experiments respectively, two groups, microinjection of DSP-4 or vehicle without IP injection of atomoxetine, will be counted the TH+ cells (n=3, n=5). TH+ cells were counted in 5 sections (n=5) for each animal, and compare DSP-4 treated with vehicle treated mice(N=3).

### 2.4 ECG recordings

Each mice were placed in plexiglass chamber retainer with body restraining before ECG monitoring (ECG-2303B, Guangzhou 3Ray Electronics Co., Ltd). Connecting the mouse limbs to the clips of electrocardiographic lead wires in sequence. After the connection was completed, the ECG recording was performed after the mice have adapted. DBA/1 mice from the different pre-treatment groups, the monitoring of ECG was performed before and post acoustic stimulation or IP administration of PTZ. The ECG experimental groups for esmolol injection were divided into 2 models as described above, the time of ECG recording started prior to IP atomoxetine (15mg/kg) for 10 min and after acoustic stimulation for 10 min or after IP PTZ (75mg/kg) for 1h. The ECG recorded at 25mm/s speed and 10mm/mv sensitivity. Arrhythmia analysis based on ECG features in human and mice^24–26^.

### 2.5 In vivo fiber photometry

#### 2.5.1 Stereotactic surgery

DBA/1 mice at 8 weeks of age were anesthetized with 3.5% chloral hydrate and head-fixed in a stereotaxic apparatus (68018, RWD Life Science Inc., Shenzhen, China), as previously described^20^. Virus (100nl) was respectively injected into the bilateral LC(AP±0.9mm; ML - 5.45 mm; DV - 3.65 mm) based on the mouse brain atlas at the rate of 40 nl/min with a 10ul microliter syringes controlled by Ultra Micro Pump (160494 F10E, WPI), the syringe was not removed until 10min after the end of infusion to allow diffusion of the viruses. Then, the optical fiber (FOC-W-1.25-200-0.37-4.0, Inper, Hangzhou, China) was implanted at the area (AP ± 0.9mmmm, ML - 5.45mm, DV - 3.55mm). The virus was allowed to express for at least 3 weeks.

#### 2.5.2 Virus

For in vivo fiber photometry, 100nl of rAAV-DBH-GCaMP6m-WPRE-hGH pAwas respectively injected into the bilateral LC of DBA/1 mice. rAAV-DBH-GCaMP6m-WPRE-hGH pA (viral titers≥ 2.00 E+12vg/ml) were purchased from Brain VTA Technology Co.Ltd (Wuhan, China).

#### 2.5.3 Fiber photometry

After 3 weeks, waiting for virus expression, postoperative mice will be used to divided into 4 groups for atomoxetine (15mg/kg, IP) or vehicle with esmolol (50mg/kg,IP) or vehicle, as described above. The fiber photometry system (Inper, Hangzhou, China, C11946) used a 488 nm diode laser. The recording started 10 min prior to IP injection of PTZ (75mg/kg) and ended after PTZ injection 1h(n=3). We analyzed data based on the events of individual trials of clonic seizures and tonic seizures derived heatmap and the values of fluorescence change (ΔF/F) by calculating (F − F0)/F0.

### 2.6 Immunohistochemistry and counting of NE neurons in LC

Mice were sacrificed and perfused with PBS and 4% PFA. After saturated in 30% sucrose(24h), each brain was sectioned into 35μm thickness of coronal slices with freezing microtome (CM30503, Leica Biosystems, Buffalo Grove, IL,USA). The brain sections were washed in phosphate-buffered saline (PBS) and then incubated in blocking solution containing 10% normal donkey serum (017-000-121, Jackson Immuno Research, West Grove, PA), 1% bovine serum albumen (A2153,Sigma-Aldrich, St. Louis, MO), 0.3% Triton X-100 in PBS for 2 hours at room temperature. For TH staining, sections were incubated at 4 ℃ overnight in rabbit anti-TH(1:1000 dilution; AB152, Merckmillipore), followed by incubation of secondary antibodies, donkey anti-rabbit Alexa 546 (1:1000; A10040, Thermo Fisher Scientific) for 2 hours at room temperature. The sections were mounted onto glass slides and incubated in DAPI (1:4000;Cat# C1002; Beyotime Biotechnology; Shanghai,China) 7mins at room temperature. Slides were sealed sheet by Anti-fluorescence attenuating tablet. All images were taken with Olympus microscope (VS120) and laser confocal microscope (Nikon A1). Stereologic cell counting of TH+ cells were performed using ImageJ software (NIH, Baltimore, MD). Images were captured using a Nikon A1 confocal microscope through 20×objective (numerical aperture 0.75). All the images were taken at a resolution of 1024×1024 pixels at RT. TH+ cells were counted in 5 sections (n=5) for each animal, and compare DSP-4 treated with vehicle treated mice (n=3).

### 2.7 Measurement the contnet and specific enzyme activity of tyrosine hydroxylase (TH) in the whole heart and heart blood by ELISA

For measurements, the DBA/1 mice were divided into two groups (n = 6/group), which included a control group without PTZ, a S-IRA group evoked by PTZ. The mice of control group were treated under anesthesia using chloral hydrate. Blood was drawn from the heart with a syringe and left to stand for 2 hours, centrifuged for 15 mins at 1000g to obtain serum. After heart blood collection, separate the heart immediately, rinse the heart with ice-cooled PBS to remove residual blood, and then follow the ELISA kit’s instructions(YS-M195, YS-M195-1, ELISA Kit, Yan Sheng Biological Technology Co., Ltd, ShangHai, China). Measure Optical Density(O.D.) at 450nm and 630nm using ELISA Microplate reader (SynergyMx M5, iD5, Molecular Devices, America).

### 2.8 Statistical analysis

All data are presented as the mean ± SEM. No statistical methods were used to pre-determine sample size. GraphPad Prism (Graph-Pad Software Inc.) and SPSS 23 (SPSS Software Inc., USA) were used for data display and statistical analysis. The incidence of S-IRA was compared in different groups using Wilcoxon Signed Rank test, as these data are nonparametric. The data on seizure scores and the latency to AGSZs/GSZs, the duration of wild running, clonic seizures, and tonic-clonic seizures were evaluated using the one-way ANOVA tests or Mann Whitney U test or Kruskal-Wallis H test. One-way ANOVA tests was used to compare TH+ cells in the LC of DBA/1 mice with and without IP injection DSP-4. Two-way ANOVA tests was used to compare heart rate with and without IP injection esmolol. The content and specific enzyme activity of TH were evaluated using the unpaired T tests. Statistical significance was inferred if p < 0.05.

## 3. Results

### 3.1 Administration of Esmolol significantly reversed the atomoxetine-mediated suppression of S-IRA evoked by acoustic stimulation

To investigate the effects of Esmolol on the incidence of S-IRA and SUDEP medicated by atomoxetine based on our previous findings, the following experiments were performed. The incidence of S-IRA evoked by acoustic stimulation in primed DBA/1 mice was obviously reduced by atomoxetine (15 mg/kg,i.p) compared with the vehicle group(n=6, n=7, p < 0.001). There was no significant difference between the vehicle group and the group pre-treated with esmolol (50 mg/kg,i.p) and with vehicle (n=7,n=6, p > 0.05). However, the incidence of S-IRA in the group pre-treated with atomoxetine (15 mg/kg,i.p) and esmolol(50 mg/kg,i.p) was significantly increased than the group pre-treated with atomoxetine(15 mg/kg,i.p) and the vehicle(n=6, n=7, p<0.05). The incidence of S-IRA in the group pre-treated with atomoxetine (15 mg/kg,i.p) and the vehicle was significantly decreased than in the group pre-treated with the vehicle and with Esmolol(50 mg/kg,i.p) (n=6, n=6, p < 0.001)(FIG 1B). The difference of latencies to AGSZs, durations of wild running, clonic seizures, tonic-clonic seizures, and seizure scores in the treatment group showed administration of esmolol significantly reversed the atomoxetine-mediated suppression of S-IRA evoked by acoustic stimulation without influencing the sensitivity to seizures (n=8, n=6, n=5, n=7, p > 0.05) (FIG 1C-F). These data showed the atomoxetine-mediated suppression of S-IRA evoked by acoustic stimulation can be significantly reversed by esmolol, which means that administration of atomoxetine reduced the incidence of S-IRA by targeting the β1-AR locating in the cardiomyocytes in our models. (FIG 1)

**Figure 1.**
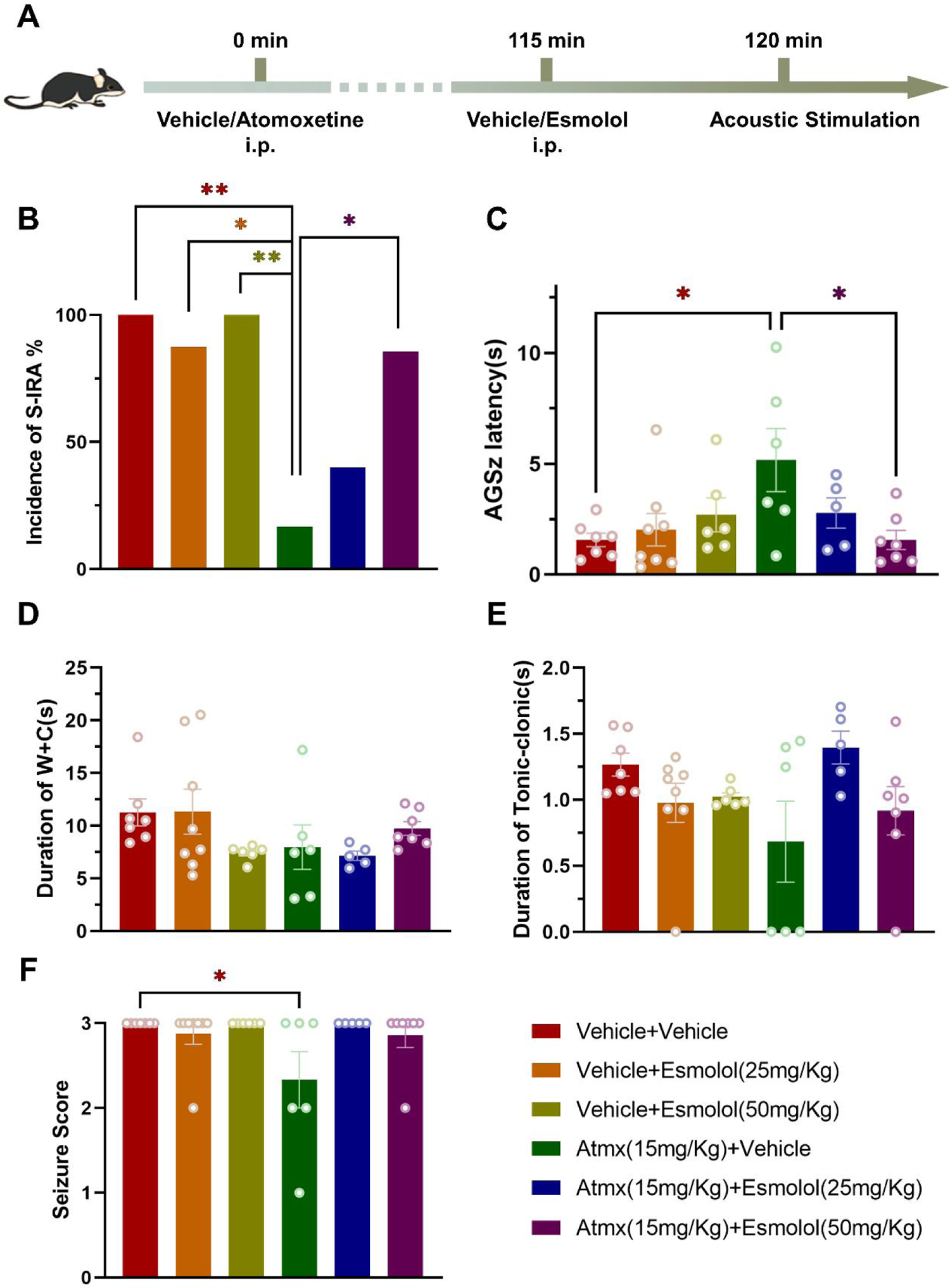
Administration of esmolol significantly reversed the atomoxetine mediated suppression of S-IRA evoked by acoustic stimulation. A. The protocol to explore the influence of administration of esmolol on atomoxetine-mediated reduction of S-IRA evoked by acoustic stimulation. B. Compared to the group treated with vehicle (n = 7), vehicle and 25mg/kg esmolol (n = 8) or vehicle and 50mg/kg esmolol (n = 6), S-IRA evoked by acoustic stimulation was markedly lower in groups treated with i.p. atomoxetine (n = 6, p < 0.05, p < 0.01) 15mg/Kg.However, the protective effect ofatomoxetine in primed DBA/1 mice was significantly reversed by esmolol doses of 50mg/kg (n = 7, p < 0.05). C. Furthermore, compared with the group treated withatomoxetine and vehicle (n = 6), the latencies to AGSZs in the control group (n = 7)or the group treated with atomoxetine and esmolol (n = 7) was significantly increased(p < 0.05). D-E. There were no intergroup differences in durations of wild running plus clonic seizures (W+C), tonic-clonic seizures (p > 0.05). F. Compared to the control group (n = 7), the seizure score was lower in the group treated with atomoxetine and vehicle (n = 6, p < 0.05). Atmx = Atomoxetine

### 3.2 Administration of Esmolol specifically reversed the atomoxetine-mediated suppression of S-IRA by targeting the β1-AR locating in the cardiomyocytes

To exclude whether the side effect of Esmolol (50 mg/kg,i.p) produces the mortality of DBA/1 mice to affect the specificity to reverse the reversed atomoxetine-mediated suppression of S-IRA in our models, the following experiments were performed in different groups. There was no significant difference in the mortality of DBA/1 mice between the group pre-treated with the vehicle and the group pre-treated with Esmolol (50 mg/kg,i.p), suggesting that the atomoxetine-mediated suppression of S-IRA can be specifically reversed by esmolol by targeting the β1-AR locating in the cardiomyocytes (FIG 2B). Compared with the group pre-treated with the vehicle, there is a transient bradycardia occurs within 10 minutes after the administration of esmolol (50 mg/kg,i.p) in the experimental group. As shown in Figure 2C, the HR was decreased most significantly at 5 minutes after the administration of esmolol (50 mg/kg,i.p). Although HR decreased significantly in the experimental group, other malignant arrhythmias such as atrioventricular block were not observed (FIG 2D-E).

**Figure 2.**
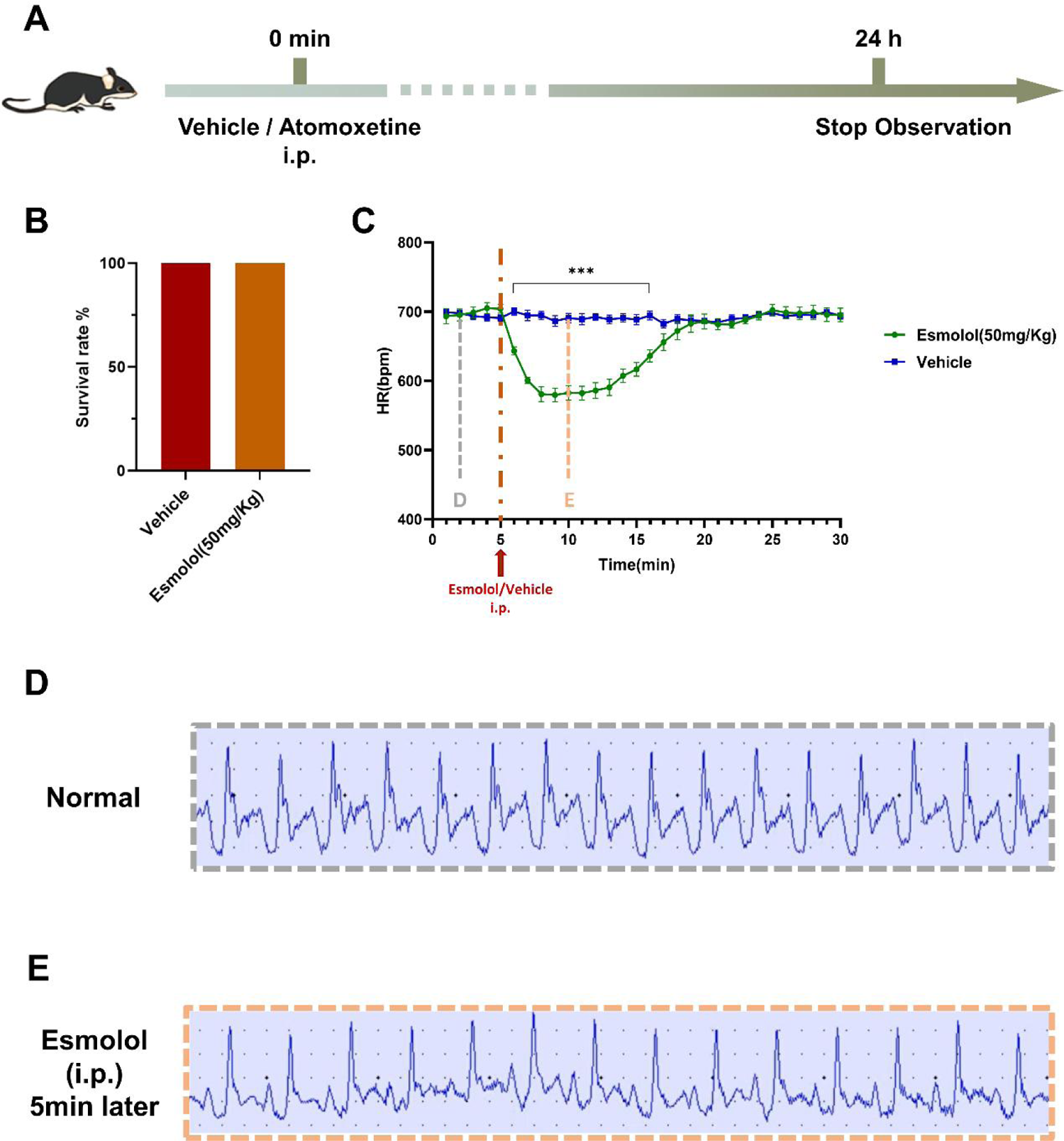
The dose of Esmolol (50 mg/kg, i.p.) does not produce the mortality of DBA/1 mice. A. The protocol to explore the dose of esmolol (50 mg/kg,i.p.) to varify whether produce the mortality of DBA/1 mice. B. There was no significant difference in the mortality of DBA/1 mice between the group pre-treated with the ehicle and the group pre-treated with esmolol(50 mg/kg, i.p, n=7) (p > 0.05). C. Compared with the vehicle group, the heart rate of DBA/1 mice in the group pre-treated with esmolol significantly decreased at 6-16 mins(p < 0.001). However, no ventricular tachycardia and atrioventricular block to be found in the DBA/1 mice from the vehicle group and the group pre-treated with esmolol. D-E. Compared with the group pre-treated with the vehicle, the heart rate of DBA/1 was decreased most significantly at 5 minutes after the administration of esmolol (50mg/kg, i.p).

### 3.3 Obvious cardiac arrhythmia immediately occurred with S-IRA evoked by both acoustic stimulation and PTZ

To observe the changing of ECG in the different S-IRA model and different treatment groups to analyze the role of atomoxetine and/or esmolol in cardiac electrophysiology, we observed that the ventricular tachycardia, ventricular premature beat, and atrioventricular block frequently occurred immediately following S-IRA evoked by PTZ or acoustic stimulation in DBA/1 mice in a group pre-treated with the vehicle and with the vehicle. However, compared with the vehicle group, the arrhythmia was significantly decreased in the group pre-treated with atomoxetine (15 mg/kg,i.p) and vehicle. We also observed the arrhythmia frequently occurred in the group pre-treated with esmolol and the vehicle and the group pre-treated with atomoxetine (15 mg/kg,i.p) and esmolol, suggesting that Esmolol can specifically reverse the atomoxetine-mediated suppression of S-IRA by targeting the β1-AR locating in the cardiomyocytes without resulting in extra arrhythmia. (FIG 3-4) (FIG 6-7)

**Figure 3.**
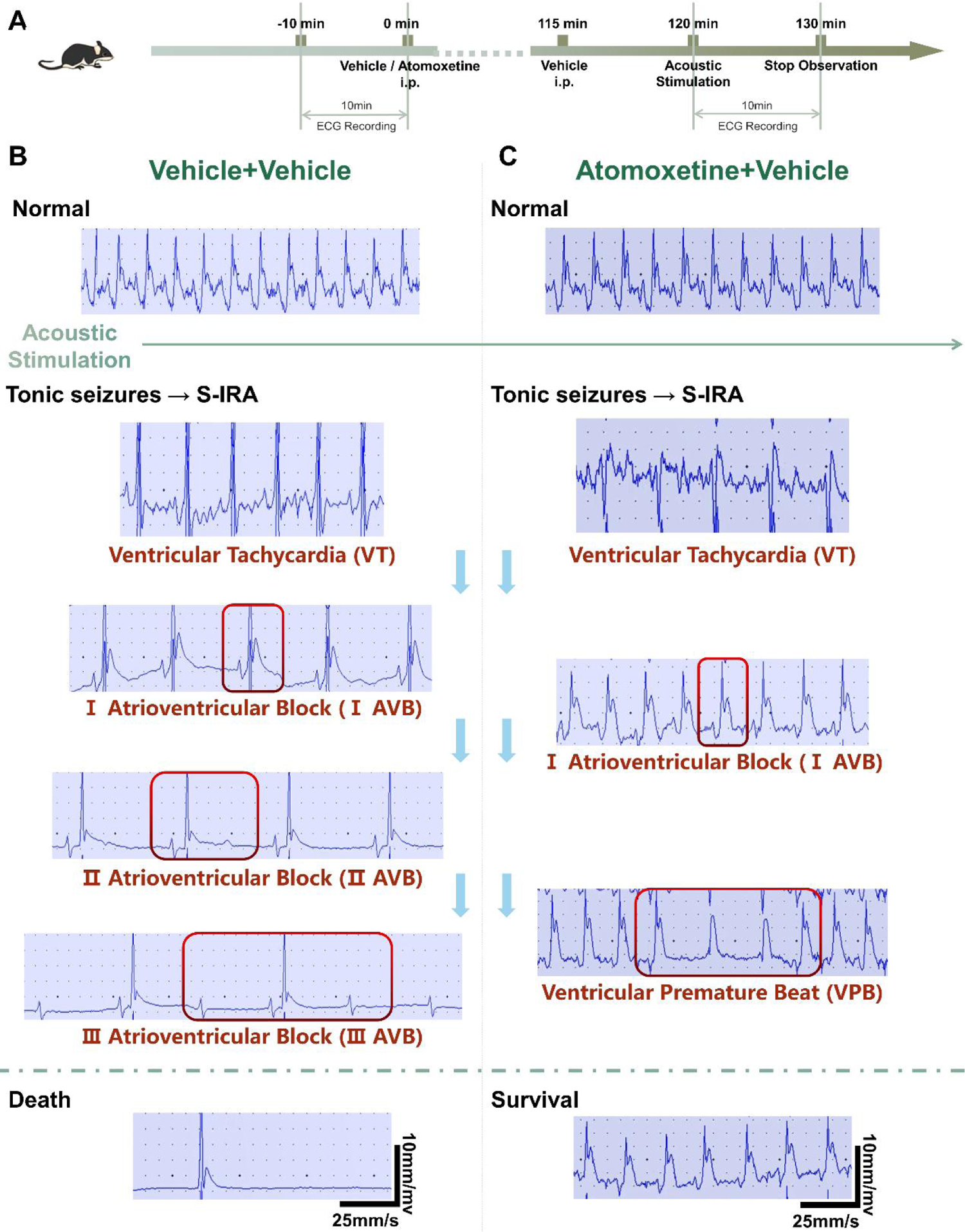
After administration of atomoxetine can obviously block the cardiac arrhythmia post S-IRA evoked by acoustic stimulation. A. The protocol to explore the changes of electrocardiogram on atomoxetine-mediated reduction of S-IRA evoked by acoustic stimulation. B. The ventricular tachycardia and atrioventricular block (I-III AVB) occurred immediately following S-IRA evoked by acoustic stimulation in DBA/1 mice in the control group. C. What’s more, the ventricular tachycardia, atrioventricular block (I AVB), and ventricular premature beat after acoustic stimulation in the group pre-treated with the atomoxetine (15mg/Kg) was observed, and the DBA/1 mice were then recovered.

**Figure 4.**
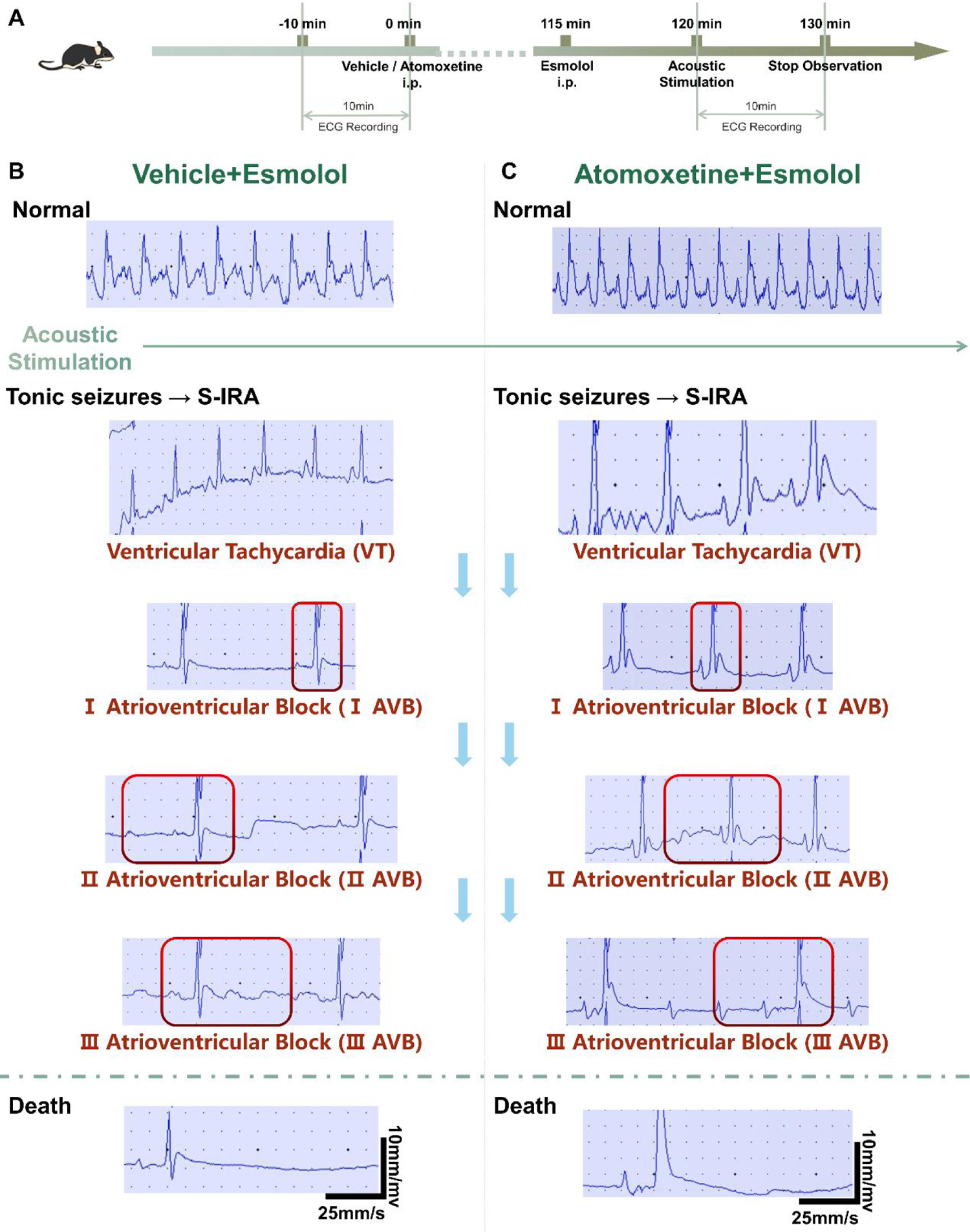
Administration of esmolol can obviously reversed the atomoxetine-mediated suppression of S-IRA evoked by acoustic stimulation via targeting the β1-AR locating in the cardiomyocytes judging by ECG analysis. A. The protocol to explore the changes of electrocardiogram on the administration of esmolol reversed the atomoxetine-mediated reduction of S-IRA evoked by acoustic stimulation. B. The ventricular tachycardia and atrioventricular block (I-III AVB) occurred immediately following S-IRA evoked by acoustic stimulation in DBA/1mice in the group treated with vehicle and esmolol (50mg/Kg). C. The ventricular tachycardia and atrioventricular block (I-III AVB) after acoustic stimulation occurred to result in the death of DBA/1 mice in the group pre-treated with the atomoxetine (15mg/Kg) and esmolol (50mg/Kg).

**Figure 5.**
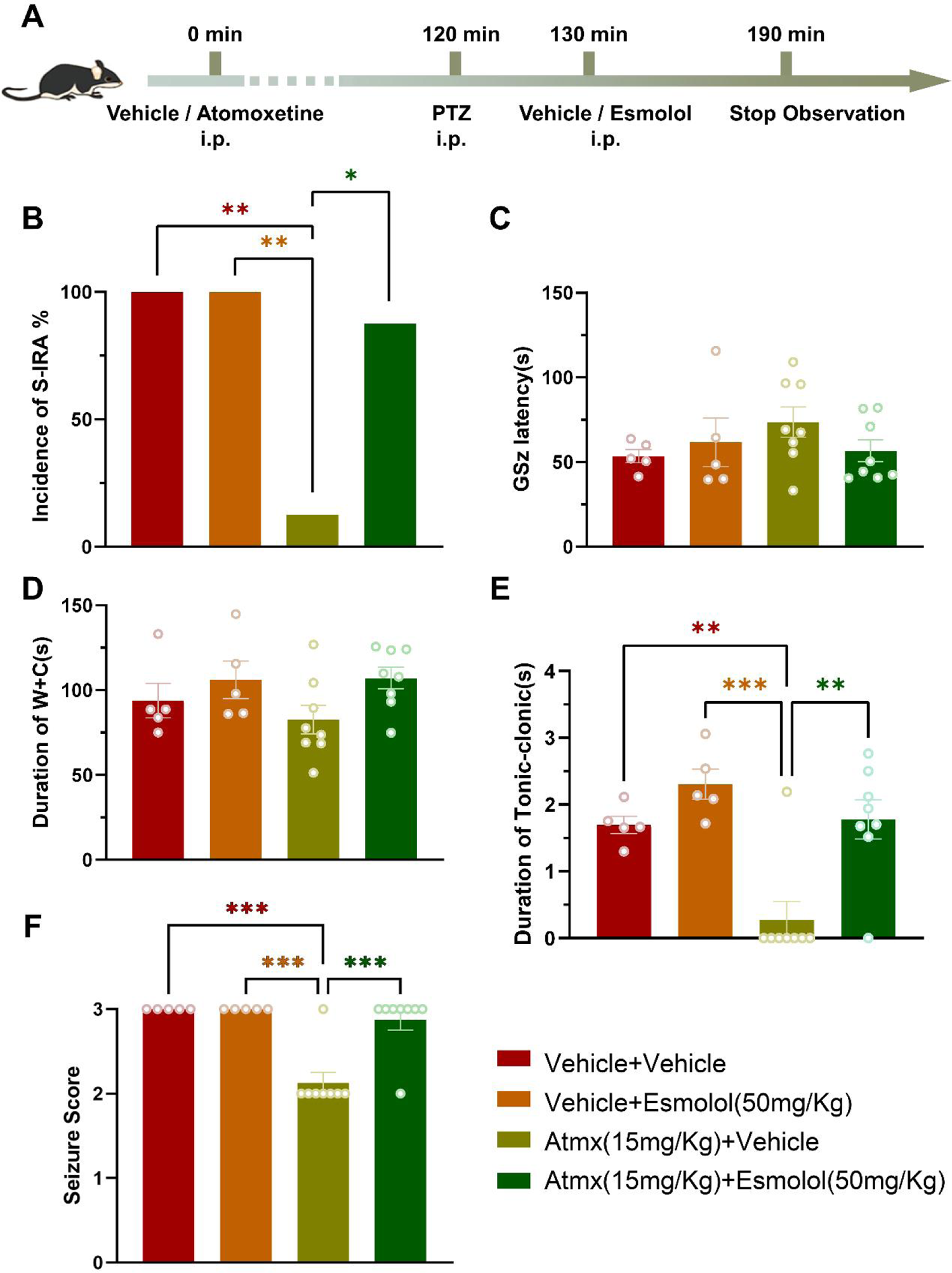
Administration of esmolol significantly reversed the atomoxetine-mediated suppression of S-IRA evoked by PTZ The protocol to explore the influence of administration of esmolol on atomoxetine-mediated reduction of S-IRA evoked by PTZ. B. Compared with the group treated with vehicle (n = 5) or vehicle and esmolol (n =5), S-IRA evoked by acoustic stimulation was markedly lower in groups treated with i.p. atomoxetine (n = 7, p < 0.01). However, compared with the group treated with i.p.atomoxetine and vehicle, the incidence of S-IRA significantly increased in the group treated with i.p. atomoxetine and esmolol (n = 7, p < 0.05). C-E. There were no intergroup differences in the GSZ latency and W+C duration (n= 6-7/group, p > 0.05). However, compared with vehicle and other groups, duration of Tonic-clonics was significantly decreased in the group with administration of atomoxetine and vehicle.F. Compared with vehicle and other groups, seizure score was significantly decreased in the group with administration of atomoxetine and vehicle.

**Figure 6.**
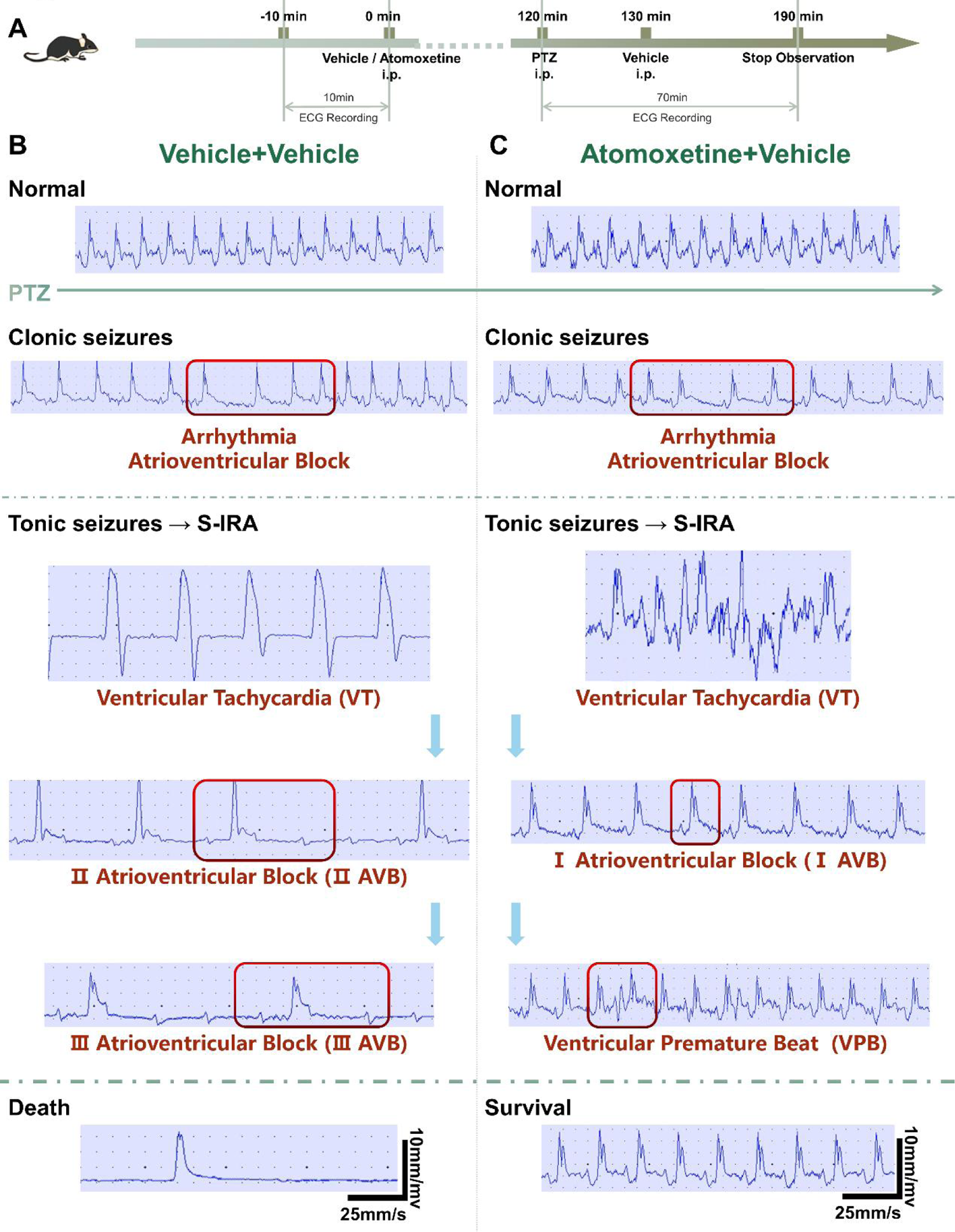
After administration of atomoxetine can obviously block the cardiac arrhythmia post the clonic seizures and tonic seizures-S-IRA evoked by PTZ. A. The protocol to explore the changes of electrocardiogram onatomoxetine-mediated reduction of S-IRA evoked by PTZ. B. The ventricular tachycardia and atrioventricular block (II-III AVB) occurred immediately following clonic seizures and tonic seizures evoked by PTZ in DBA/1 mice in the control group. C. The ventricular tachycardia, atrioventricular block (I AVB), and ventricular premature beat after PTZ injection occurred following clonic seizures and tonic seizures in the group treated with atomoxetine and vehicle, and the DBA/1 mice were then recovered.

**Figure 7.**
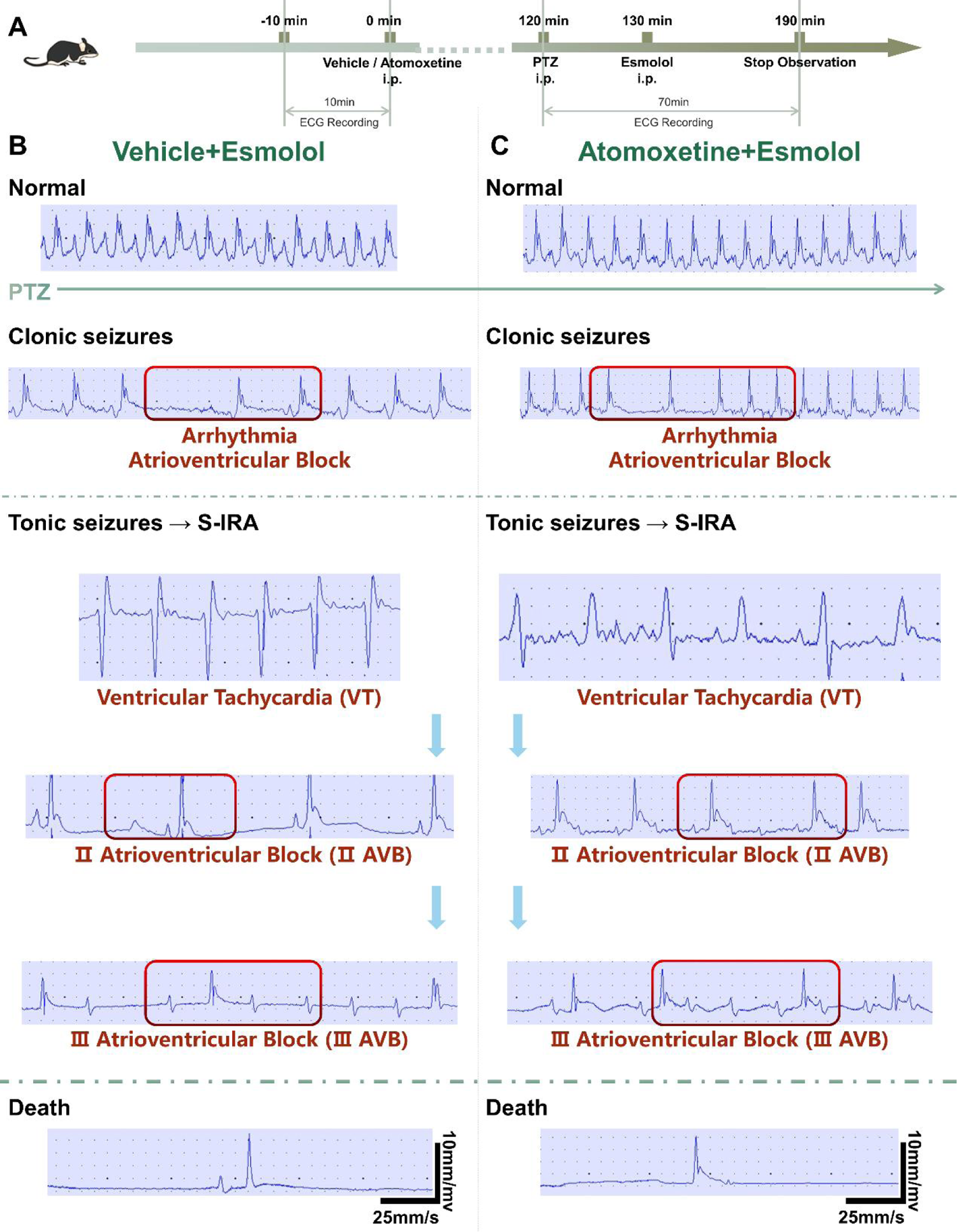
Administration of esmolol can obviously reversed the atomoxetine-mediated suppression of S-IRA evoked by PTZ via targeting the β1-AR locating in the cardiomyocytes judging by ECG analysis. A. The protocol to explore the changes of ECG on atomoxetine-mediated reduction of S-IRA evoked by PTZ. B. The ventricular tachycardia and atrioventricular block (II-III AVB) occurred immediately following clonic seizures and tonic seizures evoked by PTZ in DBA/1 mice in the control group. C. The ventricular tachycardia and atrioventricular block (II-III AVB) after PTZ injection occurred to result in the death of DBA/1 mice in the group treated with the atomoxetine (15mg/Kg) and esmolol (50mg/Kg).

### 3.4 Administration of Esmolol significantly reversed the atomoxetine-mediated suppression of S-IRA evoked by PTZ

To further investigate whether administration of Esmolol significantly reversed the atomoxetine-mediated suppression of S-IRA depends on S-IRA and SUDEP models or not, The PTZ-induced S-IRA was accepted to test in the present study. Compared with the vehicle group, the lower incidence of S-IRA by PTZ was reversed by atomoxetine (p<0.01). No obvious difference in the group of vehicle and in the group of Esmolol (p>0.05), meaning that administration of Esmolol (50 mg/kg,i.p) produced no effects on the incidence of mortality caused by PTZ itself. However, Compared with the group treated with atomoxetine and vehicle, the incidence of S-IRA and SUDEP significantly increased in the group treated with atomoxetine and Esmolol (p<0.01), suggesting that administration of Esmolol can specifically reverse the atomoxetine-mediated suppression of S-IRA evoked by PTZ. Thus, it can be speculated from our findings that enhancing the norepinephrinergic neurotransmission contributes to inhibition of S-IRA via targeting the (β1-AR) localized in the cardiomyocytes in the DBA/1 mouse SUDEP model. (FIG 5)

### 3.5 Administration of DSP-4 significantly reversed the atomoxetine-mediated suppression of S-IRA evoked by acoustic stimulation or by PTZ

Given that our previous study has showed enhancing the content of NE by the administration of atomoxetine (15 mg/kg,i.p) could significantly reduce the incidence of SUDEP in DBA/1 mice, the DSP-4 experiments were performed. According to the former study, it found that seven noradrenergic neuron populations have been identified in the CNS. Among these, neurons from LC, are the main neurons for the synthesis and release of central adrenergic neurotransmitter and have promoting effects on respiratory and cardiac networks. To examine the contributions of NE from LC neurons, we employed the DSP-4 neurotoxin, a small-molecule toxin that targets the noradrenergic transporter and selectively destroys terminals originating from LC, and in some cases cell bodies in the LC. Compared with the vehicle control group, the DSP-4 group presented dose-dependently reduced the number of TH+ cells in the LC, with the 7d post-injection group having the greatest effect (Figure 4A-C). (FIG 9). The incidence of S-IRA evoked by acoustic stimulation in primed DBA/1 mice was obviously reduced by atomoxetine(15 mg/kg,i.p) compared with the vehicle group(n=9, n=10, p < 0.05). However, the incidence of S-IRA in the group pre-treated with DSP-4 (50mg/kg, i.p) 3d or 7d before acoustic stimulation and vehicle was significantly increased than the group pre-treated with vehicle and atomoxetine(15 mg/kg,i.p) (n=6, n=5, p < 0.05). We observed the same results in the group pre-treated with DSP-4 (50mg/kg, i.p) 3d or 7d before acoustic stimulation and atomoxetine(15 mg/kg,i.p), indicating increasing the central content of NE can reduce the incidence of S-IRA(n=7, n=6, p < 0.05).(FIG 8B) Neither DSP-4 at any injection time points nor atomoxetine had a significant effect on latencies to AGSZs, durations of wild running, clonic seizures, and tonic-clonic seizures compared to vehicle group (FIG 8 C-D). The seizure scores in Fig 8F suggest that the group pre-treated with vehicle and atomoxetine(15 mg/kg,i.p) showed significantly lower seizure scores than the following pre-treated group: vehicle and vehicle(n=9, p < 0.05), DSP-4(1d) and vehicle(n=8, p < 0.01), DSP-4(3d) and vehicle(n=6, p < 0.05), DSP-4(7d) and vehicle(n=5, p < 0.01), DSP-4(1d) and atomoxetine(15 mg/kg,i.p) (n=6, p < 0.05), DSP-4(3d) and atomoxetine(15 mg/kg,i.p) (n=7, p < 0.01), DSP-4(7d) and atomoxetine(15 mg/kg,i.p) (n=6, p < 0.01) (FIG 8F). These data showed the atomoxetine-mediated suppression of S-IRA evoked by acoustic stimulation can be significantly reversed by DSP-4, which indicates that the content of NE in LC may involves in the mediating of S-IRA in our model. In the PTZ-induced SUDEP models, the atomoxetine medicated-suppressant effects of S-IRA evoked by PTZ was significantly reversed by microinjection of DSP-4 (200nl, 10ug/ul) into the every unilateral LC 7 days prior to the DBA/1 to be injected by PTZ to induce the S-IRA and SUDEP (n=8-9/group, p < 0.01). Compared with the control group, the number of TH^+^ cells in the LC in the group with microinjection of DSP-4 into LC was significantly reduced (n=8-9/group, p 7 < 0.01). (Fig10-11).

**Figure 8.**
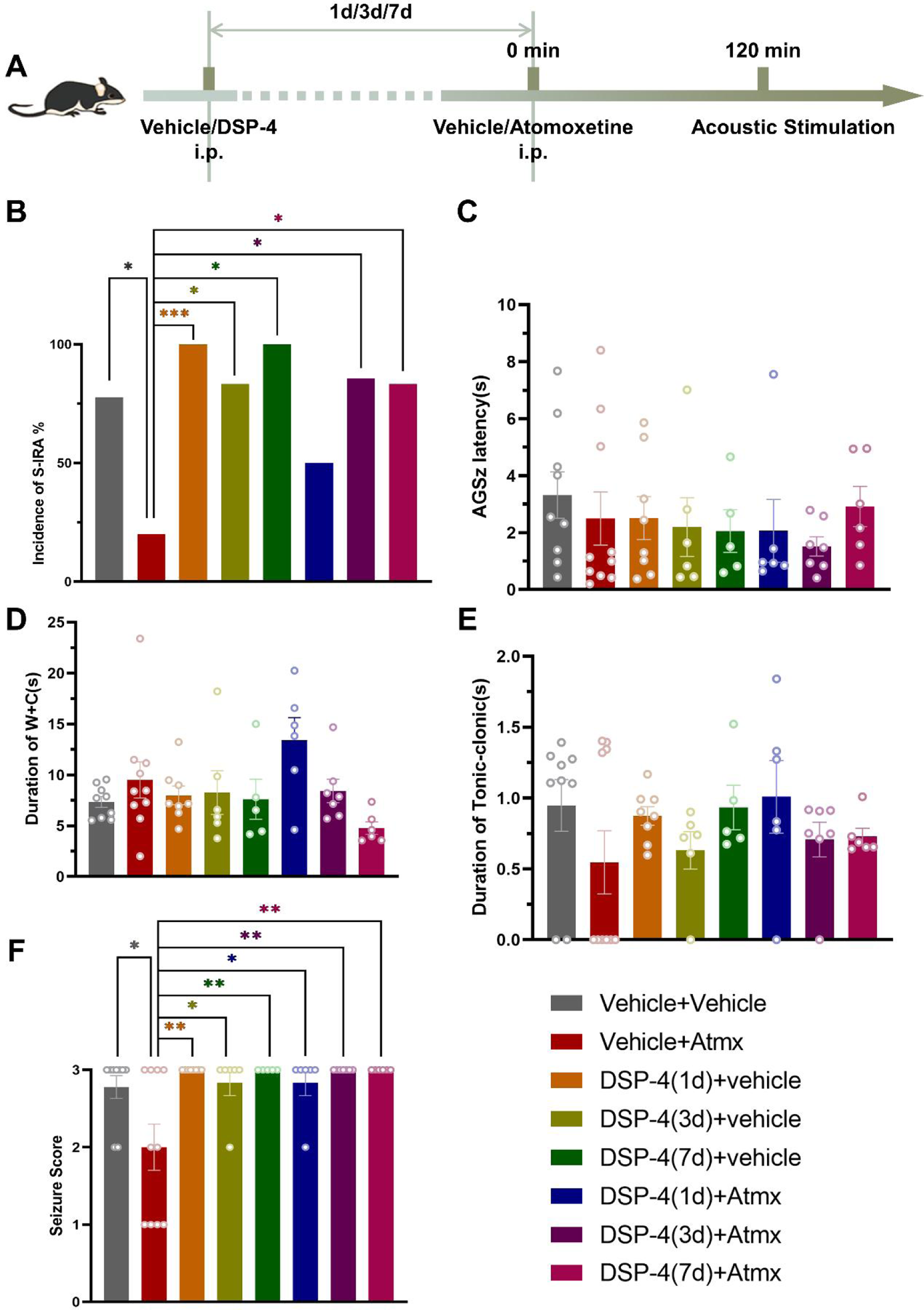
Administration of DSP-4 significantly reversed the atomoxetine-mediated suppression of S-IRA evoked by acoustic stimulation. A. The protocol to explore the influence of administration of DSP-4 on atomoxetine-mediated reduction of S-IRA evoked by acoustic stimulation. B. Compared to the group treated with vehicle and atomoxetine(n=10), S-IRA evoked by acoustic stimulation was markedly higher in groups pre-treated with i.p. DSP-4 (n = 5-8, p < 0.05,p < 0.001) and control group (n = 9, p < 0.05).However, the protective effect of atomoxetine in primed DBA/1 mice was significantly reversed by pre-treated DSP-4 3days and 7 days (n = 6-7, p < 0.05). C-E. There were no intergroup differences in the AGSZs latency, durations of wild running and clonic seizures (W+C), and durations of tonic-clonic seizures (p > 0.05). F. Compared to the group treated with vehicle andatomoxetine (n=10), the seizure score was higher in the other groups (n = 5-9, p < 0.05,p < 0.01).

**Figure 9.**
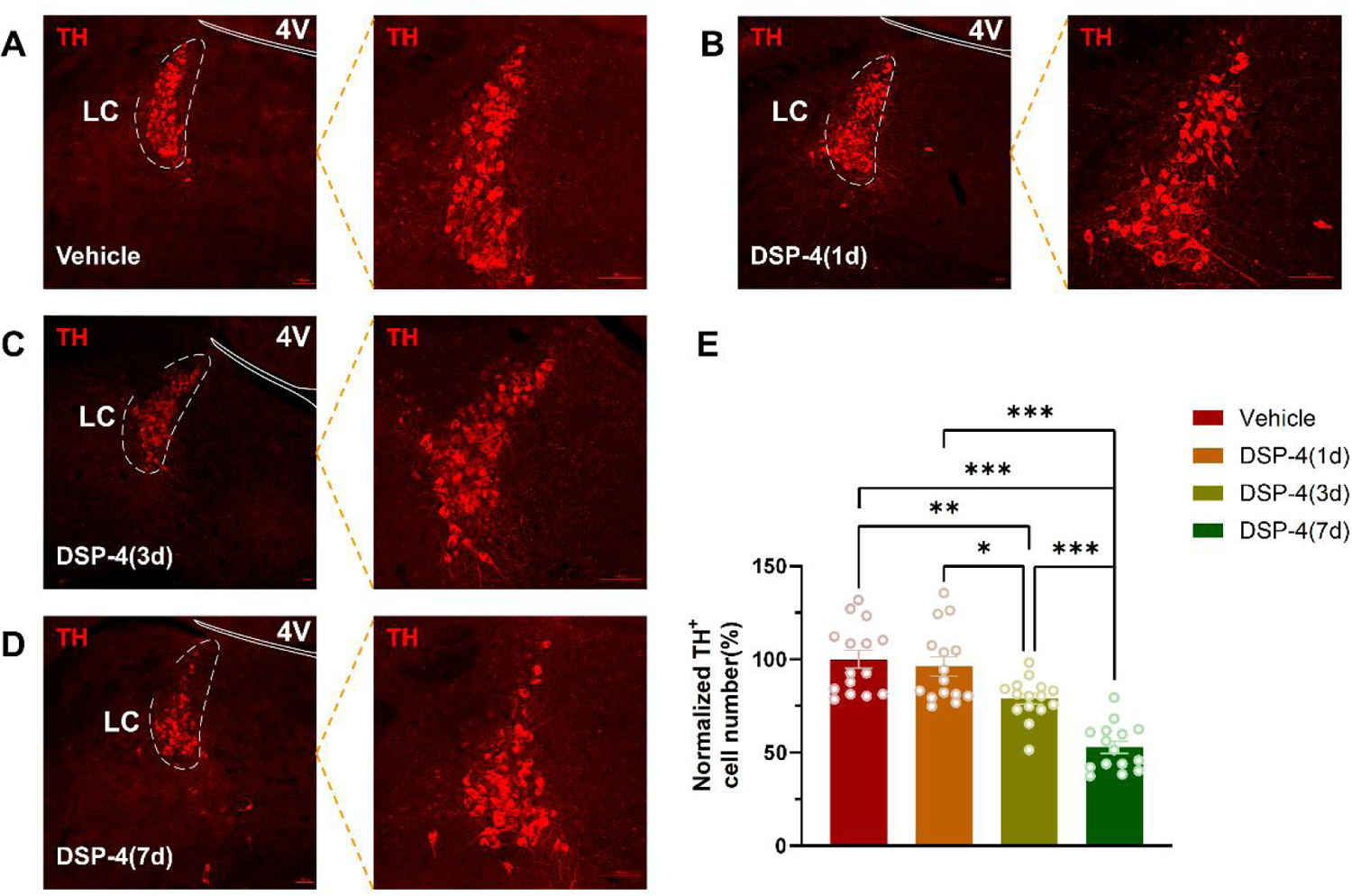
Noradrenergic neurons in LC was significantly reduced by adminstration of DSP-4. A-D. Changes of Th+ neurons in LC after intraperitoneal injection of DSP-4 or vehicle pre-treated mice for 1, 3, 7 days. E.Compared with the control group, the DSP-4 groups presented time-dependently reduced the number of TH+ cells in the LC, with the 7days post-injection group having the greatest effect (N=3, n=5,p < 0.05, p < 0.01, p < 0.001). LC: Locus coeruleus. 4V: Fourth ventricle. Confocal image magnifications:10x, 20x

**Figure 10.**
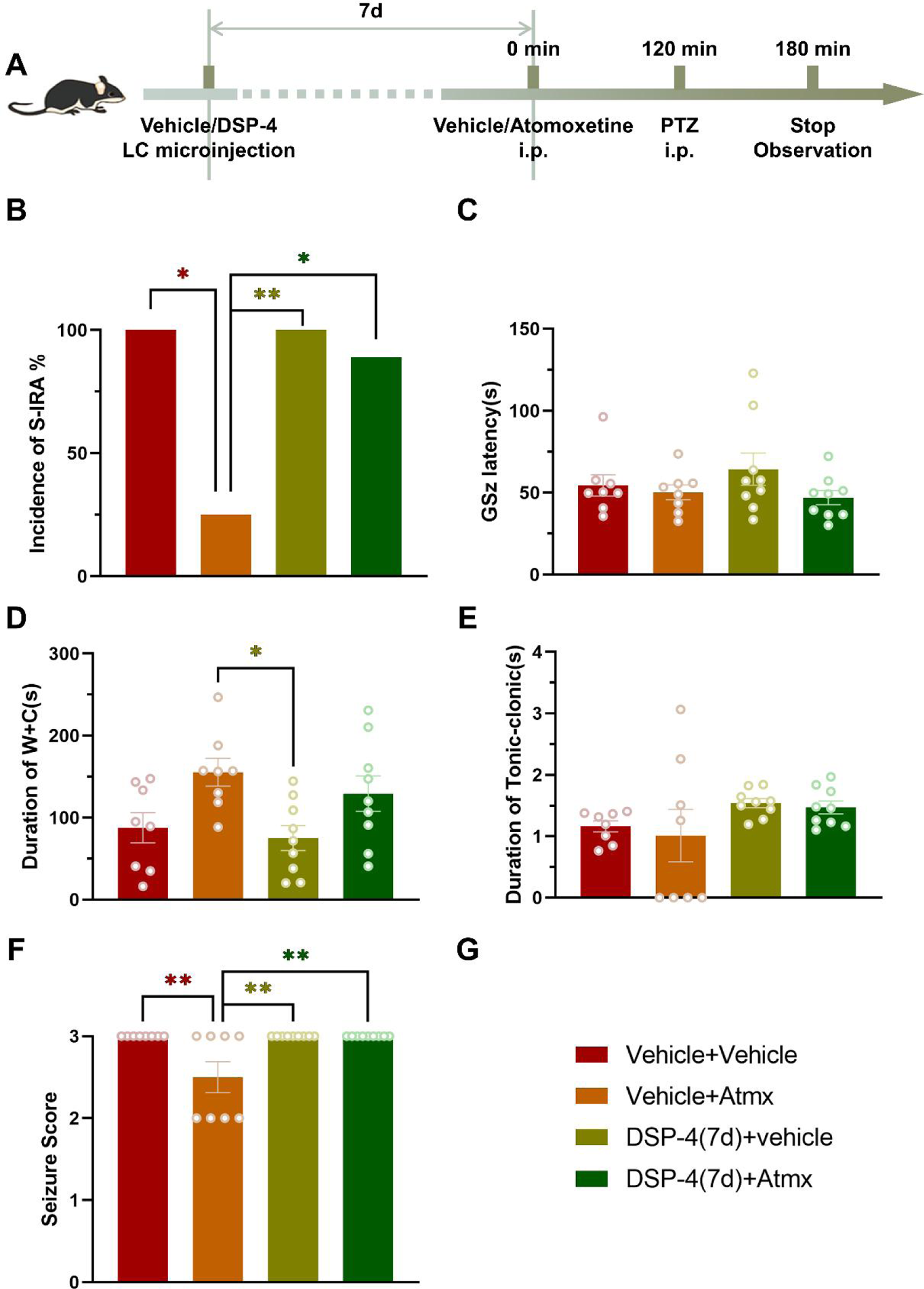
Microinjection of DSP-4 in the bilateral locus coeruleus significantly reversed the atomoxetine-mediated suppression of S-IRA evoked by PTZ. A. The protocol to explore the influence of microinjection of DSP-4 in the bilateral locus coeruleus on atomoxetine-mediated suppression of S-IRA evoked by PTZ. B. Compared with the group treated with vehicle (n = 8) or DSP-4 and vehicle (n =9), S-IRA evoked by PTZ was markedly lower in groups treated with i.p. atomoxetine (n = 8, p < 0.05,p < 0.01). However, compared with the group treated with vehicle and atomoxetine, the incidence of S-IRA significantly increased in the group treated with DSP-4 and atomoxetine (n = 9, p < 0.05). C-E. There were no intergroup differences in the AGSZs latency, durations of wild running and clonic seizures (W+C), and durations of tonic-clonic seizures (p > 0.05). F. Compared to the group treated with vehicle and atomoxetine (n=8), the seizure score was higher in the other groups (n = 8-9, p < 0.01).

**Figure 11.**
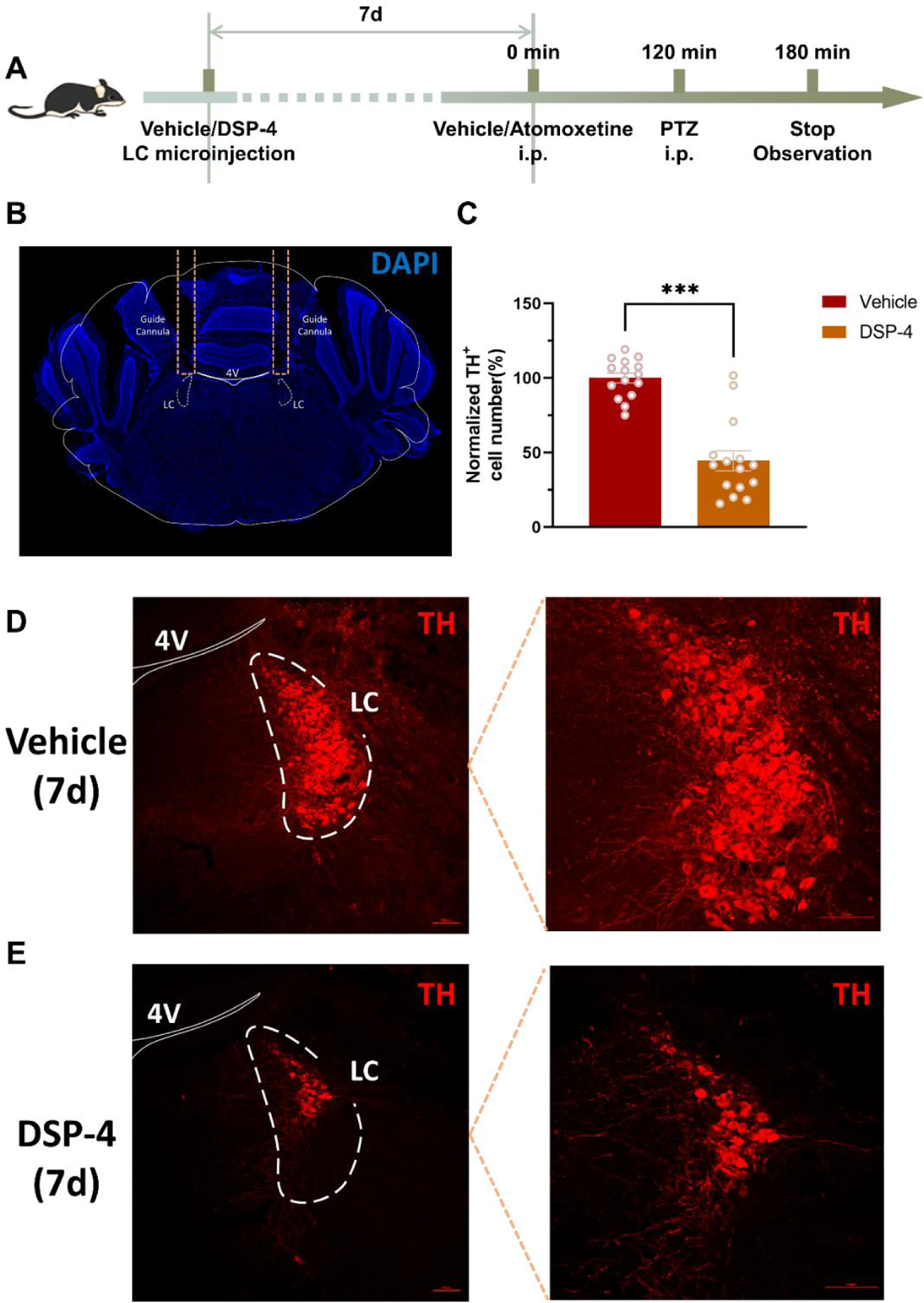
Noradrenergic neurons in LC was significantly reduced by adminstration of DSP-4 The protocol to explore the influence of microinjection of DSP-4 in the bilateral locus coeruleus on atomoxetine-mediated suppression of S-IRA evoked by PTZ. B. Representative images of implantation in DBA/1 mice for microinjection of DSP-4 into LC. C. Compared with the control group, the group pre-treated with DSP-4 through guide cannula 7 days ago and without intraperitoneal atomoxetine, TH+ cells in the LC was significantly reduced. (N=3, n=5, p < 0.001). D-E. Changes of TH+ neurons in LC after microinjection of DSP-4 or vehicle for 7 days. LC: Locus coeruleus. 4V: Fourth ventricle. Confocal image magnifications:10x, 20x

### 3.6 PTZ-induced neuronal activity of NE in LC during seizures was significantly reduced by adminstration of atomoxetine based on the photometry recordings

To investigate the activity of LC during the seizure attacking including clonic and tonic seizure phases evoked by PTZ, calcium signaling within the neurons of the bilateral LC was recorded by photometry in mice infected with GCaMP6f in the bilateral LC. The activity wave of calcium signals in the bilateral LC was strong during the phases of clonic seizures in different treatment groups. However, no obvious changing to be found in the groups with the pre-treatment of vehicle and atomoxetine, atomoxetine and Esmolol in the tonic seizures duration. These data showed that the increasing of abnormal activity within LC neurons to synthesize and release the more NE to the synaptic cleft to target its receptor postsynaptic membrane to block SUDEP. However, due to the part reuptake of NE into LC, it is insufficient to block SUDEP. Conversely, administration of atomoxetine which passes through the blood brain barrier to target the LC to block the reuptake of NE and enhance the transfer efficiency of NE within synaptic cleft to block SUDEP. (FIG 12).

**Figure 12.**
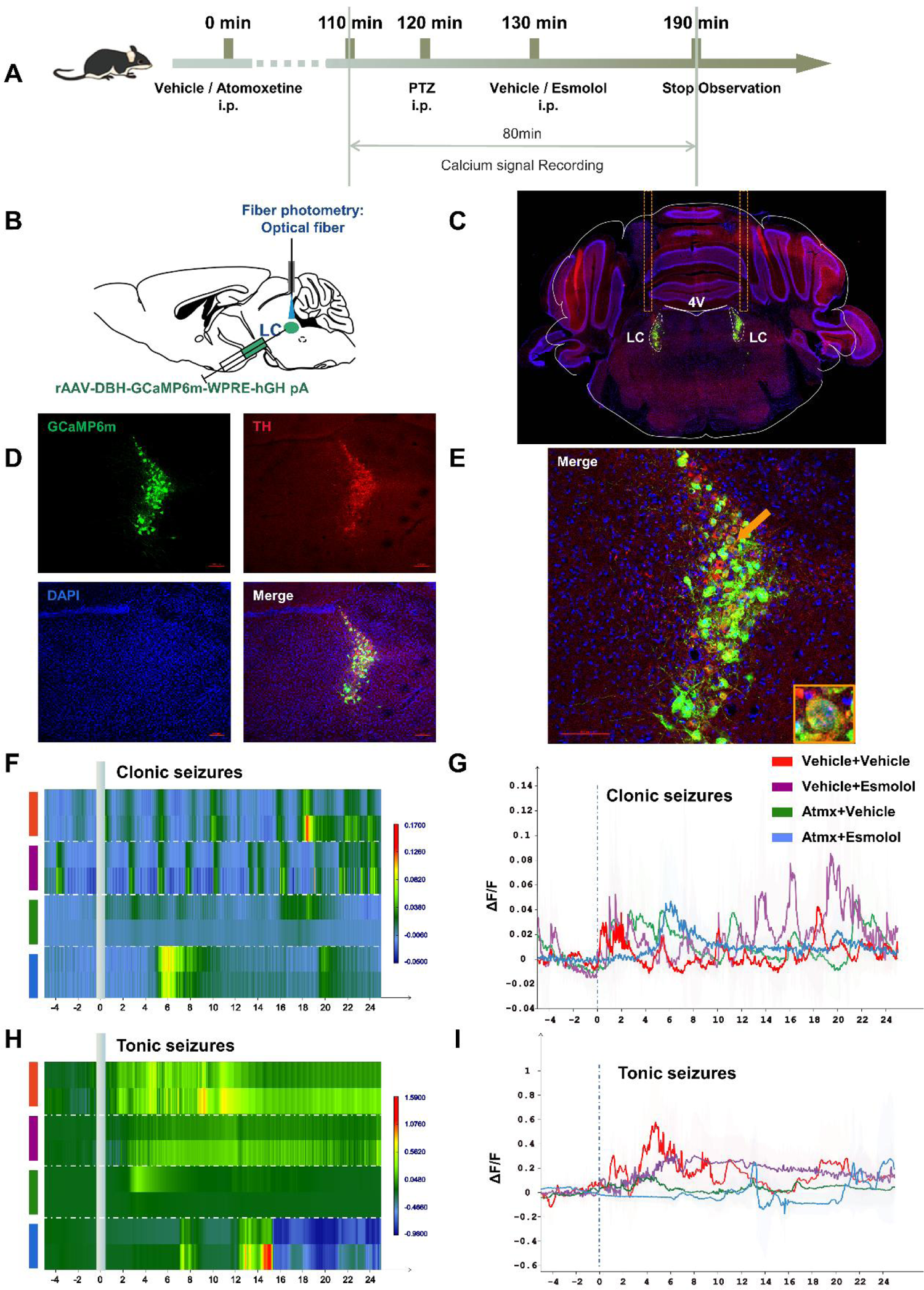
PTZ-induced neuronal activity in LC during seizures was significantly reduced by adminstration of atomoxetine based on the photometry recordings. A. The protocol to explore the influence of intraperitoneal atomoxetine on PTZ-induced neuronal activity in LC during seizures based on the photometry recordings. B-C. The position of virus injection and optical fiber D-E. Immunohistochemistry of noradrenergic neurons and Gcamp virus in LC F-I. The activity wave of calcium signals in the bilateral LC was strong during the phases of clonic and tonic seizures in the group with pre-treatment of vehicle and vehicle, vehicle and esmolol, atomoxetine and esmolol, compared with the group with the pre-treatment of atomoxetine and vehicle.

### 3.7 PTZ-induced TH activity but not TH content from the serum of left ventricle and the whole heart tissue was reduced following the S-IRA

To investigate the role of activity of NE from the sympathetic preganglionic fibers of heart to modulate SUDEP, the content and activity of TH from the serum of left ventricle and the whole heart tissue of DBA/1 mice were analyzed by ELISA, respectively. Compared with the control groups, the content of TH from the serum of left ventricle and the whole heart tissue of DBA/1 mice were significantly increased following S-IRA evoked by PTZ (n=6/group, p < 0.01). Conversely, Compared with the control groups, the activity of TH from the serum of left ventricle and the whole heart tissue of DBA/1 mice were significantly reduced following S-IRA evoked by PTZ (n=6/group, p < 0.05). The above data suggested that the lower activity of TH sympathetic preganglionic fibers of heart may play a key role to lead to S-IRA and SUDEP. (FIG 13)

**Figure 13.**
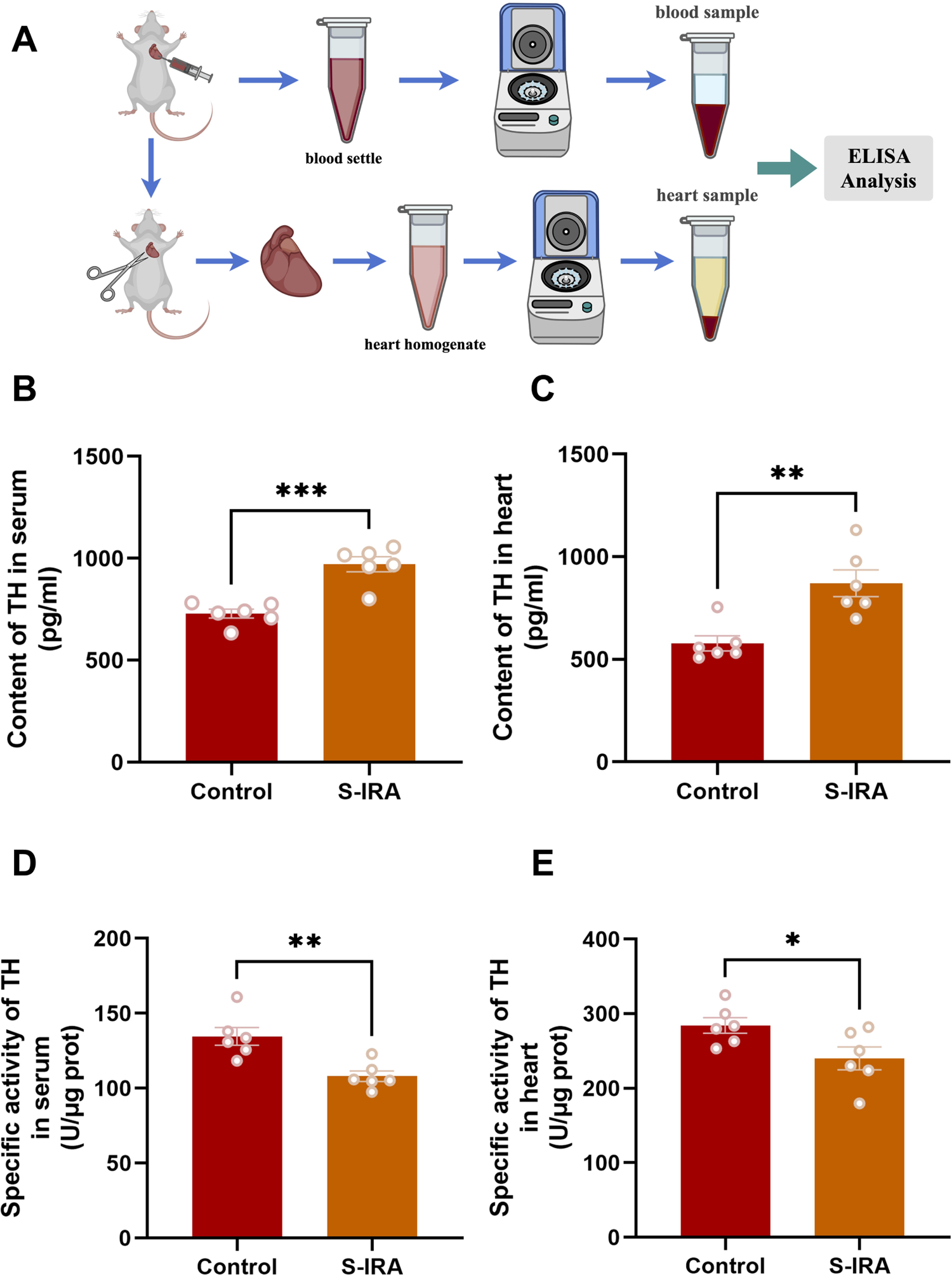
PTZ-induced TH activity but not TH content from the serum of left ventricle and the whole heart tissue was following the S-IRA, respectively. A. The protocol to investigate changing the content and activity of TH from the serum of left ventricle and the whole heart tissue of DBA/1 mice were analyzed by ELISA, respectively. B-C. Compared with the control groups, the content of TH from the serum of left ventricle and the whole heart tissue of DBA/1 mice were significantly increased following S-IRA evoked by PTZ (n=6/group, p < 0.01). D-E. Compared with the control groups, the activity of TH from the serum of left ventricle and the whole heart tissue of DBA/1 mice were significantly reduced following S-IRA evoked by PTZ (n=6/group, p < 0.01, p < 0.05).

**Figure 14.**
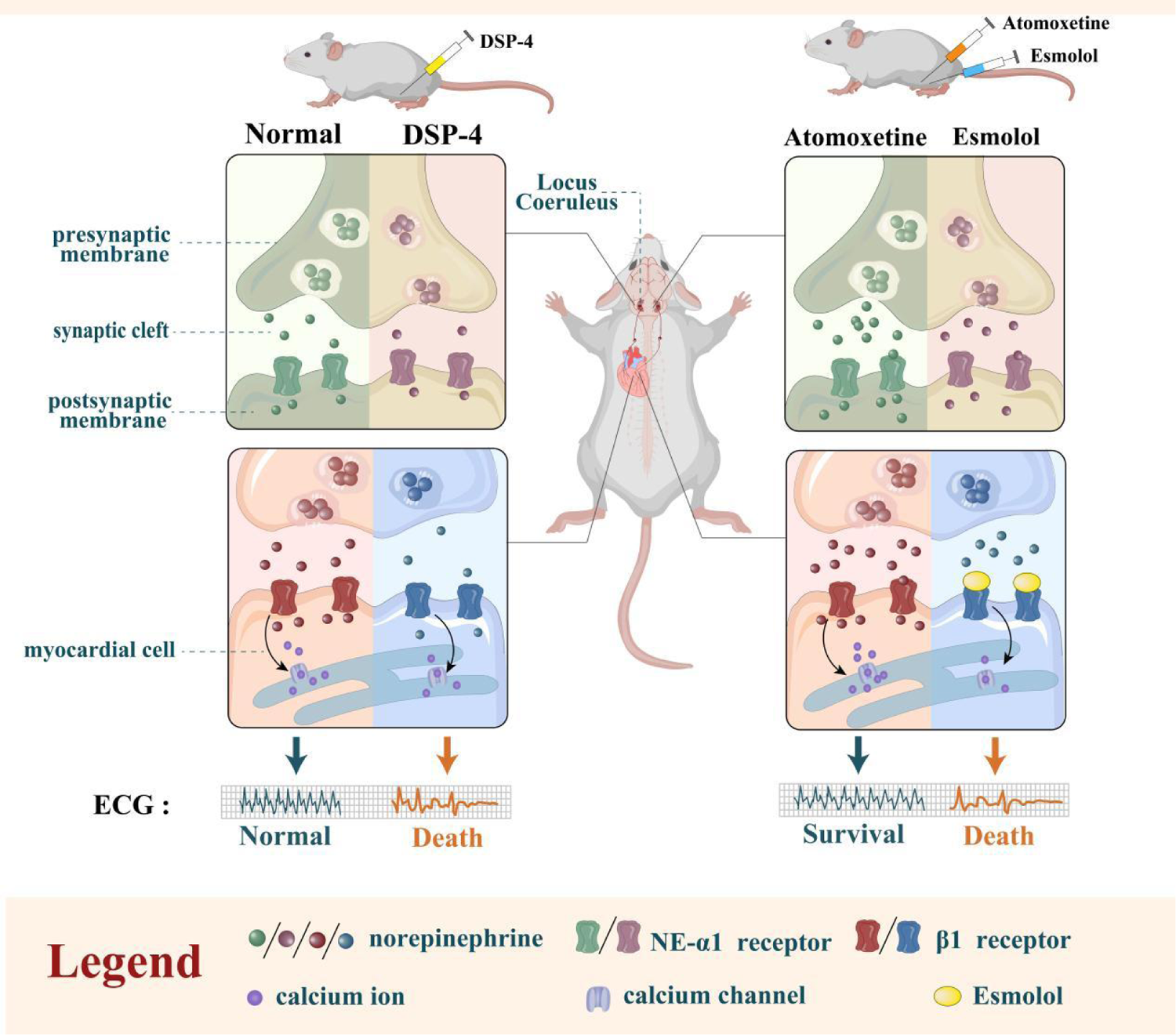
The interaction between the noradrenergic neurons in the locus coeruleus and beta-1 adrenergic receptor (β1-AR) in the cardiomyocytes was implicated in modulating SUDEP

## 4. Discussion

There are about 70 million people in the world who suffer from epilepsy, and 2.4 million have been diagnosed with epilepsy every year, nearly 90% occurred in developing countries^1^. The latest epidemiological studies of the American Epilepsy Society show that the incidence of SUDEP in western developed countries is about 1.11 per 1000 people per year^2^, while the results of the multi-center epidemiological survey in China show that the incidence of SUDEP in China is about 2.3 per 1000 people per year^4^, and the number of domestic epilepsy patients dying from SUDEP is more than 20,000 per year. More importantly, SUDEP is the main factor that leads to the death in young patients with epilepsy with an incidence two to three times that of other diseases that can lead to sudden death in young adults, and children’s SUDEP incidence is also close to adults^3^. Japan’s latest epidemiological survey shows that the incidence of SUDEP in pregnant women is significantly higher than the incidence of non-pregnant women^6^. However, due to the complexity of its pathogenesis and the difficulty to obtain the real-time monitoring data of SUDEP in patients with epilepsy, the understanding of the SUDEP mechanism is still limited, and it is difficult to develop a prevention strategy. Many previous studies have shown that the pathophysiology of SUDEP is complex, including the disfunction of heart, autonomic nerves, respiratory, brain structural obstacles, and polygenesis^5, 11^.

The previous research has shown that the main risk factors of SUDEP are generalized tonic-clonus and nocturnal seizures. The neuronal network of generalized tonic-clonic seizures may lead to hypoventilation, apnea, and cardiovascular failure by inhibiting brainstem respiration or autonomic nerve control centers, leading to death^5, 11^. In recent years, more and more researches have shown that the apnea and cardiac dysfunction are important potential mechanisms in SUDEP. Studies in animal models and SUDEP patients have shown that S-IRA is one of the main induced factors that lead to death in many cases^12, 27, 28^. Our previous findings showed that administration of atomoxetine inhibited S-IRA and SUDEP in DBA/1 mice independently of seizure induction methods, and its inhibition of S-IRA induced by acoustic stimulation was dose-dependent and the suppression of S-IRA by atomoxetine could be mediated by NE/NEα-1R interactions that can be reversed by prazosin, a NEα-1R antagonism. Given the current clinical reports that atomoxetine can produce the effects of peripheral circulation^29^. For example, after oral administration of atomoxetine, ADHD (Attention-Deficit/ Hyperactivity Disorder) patients showed symptoms such as increased heart rate and other symptoms^30^, considering that atomoxetine may play a role in SUDEP not only through the central - peripheral - cardiac sympathetic mechanisms. What’s more, the recent study showed that the tonic phase apnea evoked by acoustic stimulation is not sufficient for seizure-induced death in a W/+Emx1-Cre mouse. A respiratory arrest is not the only lethal cause, tonic-clonic seizures caused sudden, simultaneous respiratory and cardiac depression in mice, and the time of recovery of respiration and heart rate during resuscitation is also closely matched^31^. Therefore, the heart also plays an important role in the mechanism of SUDEP^32^.

Seizure-associated arrhythmias are common and have been considered as the potential pathogenesis of SUDEP, such as tachycardia, bradycardia, and cardiac arrest after the seizures, resulting in cardiac repolarization, changes in electrolyte levels, blood pH, and catecholamine release in patients with epilepsy, which latter three factors may promote arrhythmias and lead to death by affecting cardiac excitability^33^. Tachycardia and several factors that increase susceptibility to tachyarrhythmias, known as pathological cardiac repolarization, are considered as risk factors for sudden cardiac death (SCD) in healthy individuals and patients with epilepsy which, in this particular setting, are important factors in triggering SUDEP events^33, 34^. However, the exact mechanism of SUDEP and cardiac arrest induced by seizures remains unclear. Seizures may interfere with the normal functioning of the heart by causing autonomic disturbances^35^, leading to fatal arrhythmias. Studies have found that cardiac electrical instability and autonomic dystonia are caused by cumulative cardiac damage during repeated attacks^36^. Studies also have shown that catecholamine (norepinephrine and adrenaline) released from seizures is related to a wide range of calcium (Ca^2+^)-mediated physiological changes, which can lead to damage to the heart structure. Our previous studies found that repeated S-IRA in DBA / 1 mice will result in necrotic damage in the ventricle which is caused by the local Ca^2+^ homeostasis disturbances, and the incidence and severity of injury depend on the total number of S-IRA^12^. Relevant clinical data have found that paroxysmal arrhythmias (including arrest, atrioventricular block, and rare atrial and ventricular fibrillation) usually occur after convulsive episodes and are often associated with (near) SUDEP^37^. Studies have shown that chronic epilepsy can lead to hypoxemia and the increase of catecholamine, causing damage to the heart and vascular, resulting in cardiac electrical, and mechanical dysfunction^38, 39^. Some fatal incentives may be the acidosis caused by persistent hypoxemia and hypercapnia, leading to bradycardia or cardiac arrest, while in others it may be the seizures due to malignant arrhythmias^40^. This suggests that SUDEP is inextricably linked to arrhythmia.

In the present study, due to the consideration for the β1 receptor which locates preferentially in the cardiomyocytes of the heart can mediate the Ca^2+^ steady state of cardiomyocyte endoplasmic reticulum to modulate the heart rhythm and myocardial contractility, we used the β1 receptor blocker esmolol to block the β1 receptor to explore the connection between SUDEP and heart. It turned out that the suppressant effects of S-IRA by atomoxetine were markedly reversed by esmolol, suggesting that administration of atomoxetine significantly reduced the incidence of S-IRA by enhancing the concentration of norepinephrine of the cleft between the sympathetic synapse and cardiomyocytes of the heart to combine with β1 receptors of cardiomyocytes. The ECG of DBA/1 mice with administration of esmolol (50 mg/kg, i.p) that can significantly reverse the lower incidence of S-IRA evoked by acoustic stimulation by administration of atomoxetine showed the mixture of sinus bradycardia, atrioventricular block, ventricular premature beat, and ventricular tachycardia in order. In the meantime, the ECG of DBA/1 mice suffered from the S-IRA evoked by acoustic stimulation characterized by the mixture of sinus bradycardia, atrioventricular block, ventricular premature beat, and ventricular tachycardia as well in our model. However, no obvious arrhythmia appeared in the DBA/1 mice without suffering from S-IRA other than the sinus bradycardia and no apparent mortality occurred in the group of DBA/1 mice with pre-treatment with the dose of esmolol 50 mg/kg (i.p). Furthermore, the arrhythmias can be significantly reduced by atomoxetine in our models. Thus, the lower incidence of S-IRA by acoustic stimulation can be significantly as well as especially reversed by esmolol by blocking the β1-AR located in cardiomyocytes to combine the norepinephrine released by cardiac sympathetic nerve synaptic terminal in our models.

Additionally, it is possible that the locus coeruleus (LC), as the largest nuclei to synthesize and release norepinephrine in the brain, may be implicated in the course of administration of esmolol to reverse the lower incidence of S-IRA and SUDEP by atomoxetine. We observed the activity of neurons in LC by recording calcium signals through optical fibers. Firstly, specific promoter DBH-GCaMP6m to target the TH neurons in LC was used, and it was found that neuronal calcium signaling activity increased in LC during the clonic.Conversely,, no different changing of activity of neurons to be found during the tonic seizures stage in the group pretreatment with atomoxetine.

Although the findings of fiber photometry showed that the noradrenergic neurons activity in the LC in our models, indeed, was implicated in modulating SUDEP, whether the the noradrenergic neurons in the LC directly mediate the course and how to interact with the β1 receptor needs further to be explored. Based on this consideration, we pretreated DBA/1 mice with DSP-4, a selective neurotoxin for the LC noradrenergic system to test it, which can induces acute and relatively selective degeneration of central noradrenergic nerve terminals^41^. After 3 or 7 days of administration of DSP-4 by intraperitoneal injection, DBA/1 mice were injected with atomoxetine and given acoustic stimulation. We found that administration of DSP-4 could reverse the reduction of the incidence of S-IRA by atomoxetine. By staining and counting TH-labeled neurons in the LC, it showed that after 3 or 7 days for DSP-4 treatment, the TH-labeled neurons decreased significantly. It is inferred that LC may play a key role in modulating S-IRA and SUDEP by NE neurons and regulating the respiratory and circulation function in our models. However, due that we chose to degrade the noradrenergic neurons in the LC by IP administration of DSP-4 which can degrade other regions of brain, especially including the heart and other peripheral noradrenergic neurons system. Thus, it is too difficult to verify whether the noradrenergic neurons in the LC modulate SUDEP with interacting the β1 receptor of heart. Therefore, we further select the pathway of microinjection of DSP-4 into the LC to degrade the noradrenergic neurons in our models to test the above experimental data. It turned out that the results of IP administration of DSP-4 was basically consistent with of microinjection of DSP-4 into the LC, which strongly suggesting that the noradrenergic neurons in the LC by interacting the β1 receptor of heart may a key role in modulating S-IRA and SUDEP in our models.

In the current study, we firstly reported that the suppressant effects of S-IRA and SUDEP in DBA/1 mice by atomoxetine can be significantly reversed by administration of esmolol, a selective β1 receptor blocker and connected brain and heart in SUDEP pathomechanism. Therefore, we assumed that seizure, especially in the early stage of GTCS, causes excitation of central sympathetic nerve, depleting the NE released by LC, and then reduces the binding efficiency of β1-AR on myocardial cell membrane, leading to arrhythmia and eventual SUDEP. Nevertheless, the specific mechanisms such as which nucleus or neurons are involved in the axis from LC to the heart still need to be further explored. Esmolol has been widely used in clinical practice. The study of the relationship between the LC and the β1-AR of heart to be implicated in SUDEP will provide a new target for the prevention of SUDEP and pose the clinical transformation significance.

During the research, mice simultaneously experienced S-IRA and arrhythmia, but the sequential order was not recorded. A previous study showed that S-IRA evoked by acoustic stimulation in DBA/1mice was characterized by the simultaneous suppression of respiration and circulation though the respiratory arrest preceded the cardiac arrest^31^. Clinical trials have found that most patients with sudden epilepsy experienced autonomic system failure, beginning with respiratory dysfunction, followed by heart failure. It may support the idea that brain stem structure that controls autonomic function is involved in the mechanism of SUDEP, and that there are a series of abnormal breathing patterns before heart abnormalities. Therefore, we will have further research in the sequence of suppression of respiration and circulation during SUDEP.

Nevertheless, in the present study showed that administration of atomoxetine enhancing the content of NE within synaptic cleft by blocking the reuptake of NE into the LC to prevent SUDEP. Importantly, microinjection of DSP-4 into the LC to partly degrade NE neurons can reverse the atomoxetine - mediated suppressant effects of SUDEP. Taken together, it is sufficient to confirm that LC was directly involved in modulating SUDEP by targeting the β1-AR in the cardiomyocytes in our models.

To further explore the role of the sympathetic preganglionic fibers of heart to modulate SUDEP, we examined the changing of the content and activity of TH in the heart from the serum of left ventricle and the whole heart tissue in our models, respectively. The data showed that the content was significantly increased following the S-IRA of DBA/1 mice, respectively. In contrast, the activity of TH was significantly decreased post the S-IRA of DBA/1 mice. Based on our previous results that the TH activity the low brainstem in the DBA/1 mice was increased following the S-IRA to be resuscitated by mechanical ventilation, the above data may indicate that the exhaustion sympathetic nerve and the low activity of TH in the latter stage of seizure may be a key factor leading to SUDEP. However, the exact role both of central and peripheral sympathetic nerve and TH activity still to be further explored in our models.

Nevertheless, our main findings had been firstly exhibited as follows: 1) Atomoxetine medicated the S-IRA and SUDEP by targeting the LC. 2) NE neurons in the LC directly were implicated in modulating the S-IRA and SUDEP including both of respiration and circulation factors. 3) The S-IRA was not the independent cause leading to SUDEP. Instead, the interaction between the brain and heart, especially the LC and heart will be a promising direction to decode SUDEP.

### 4.1 The limitation to the present study

First, we used a limb lead electrocardiogram machine to record the changes of the electrocardiogram only during the S-IRA and death instead of the entire experiment. Also, in the process of measurement, the electrocardiogram was unstable at the initial stage of measurement due to the body movement of mice. Therefore, we may adopt the measurement of electrocardiogram implanted in the body to avoid the influence of mice strenuous movement during ECG measurement and monitor the ECG changes throughout the whole experiment in further study. Second, we failed performed the timely observation of heart index, ejection fraction in our models. Third, although we chose DSP-4 to degrade the NE neurons in the LC make the NE neurons lose its function to test whether SUDEP can be modulating by the axis between the LC and heart, we failed to confirm the interaction between the LC and heart by the specific technology such as optogenetics. Currently, we have proposed that there is a close connection between the brain-heart axis, especially LC, and SUDEP, but the problems that by which nucleus does LC send signals down to the heart and where specific abnormal conduction pathway of heart is still need to be solved.

## 5. Conclusions

The current finding in our models suggested that enhancing central norepinephrinergic neurotransmission in the brain, especially in the LC contributes to inhibition of seizure-induced respiratory arrest and SUDEP by targeting β1-AR locating in the cardiomyocytes. The norepinephrinergic neurotransmission-β1-AR pathway between the LC and heart may be a potential and specific target to prevent SUDEP.

